# AlphaFold encodes the principles to identify high affinity peptide binders

**DOI:** 10.1101/2022.03.18.484931

**Authors:** Liwei Chang, Alberto Perez

## Abstract

Machine learning has revolutionized structural biology by solving the problem of predicting structures from sequence information. The community is pushing the limits of interpretability and application of these algorithms beyond their original objective. Already, AlphaFold’s ability to predict bound conformations for complexes has surpassed the performance of docking methods, especially for protein-peptide binding. A key question is the ability of these methods to differentiate binding affinities between several peptides that bind the same receptor. We show a novel application of AlphaFold for competitive binding of different peptides to the same receptor. For systems in which the individual structures of the peptides are well predicted, predictions in which both peptides are introduced capture the stronger binder in the bound state, and the other peptide in the unbound form. The speed and robustness of the method will be a game changer to screen large libraries of peptide sequences to prioritize for detailed experimental characterization.

## Introduction

Machine learning has revolutionized protein structure prediction, achieving an unprecedented level of accuracy that is transforming the field.^1,2^ Deep neural network hallucination^3^ has already served to design novel proteins based on the idealized version of proteins the network learns. The structural biology community has rapidly used these developments to test neural networks beyond their original application to problems including alternative conformational sampling, disordered protein identification, or complex structure prediction, with very promising results^4,5^.

At the early stages of complex structure prediction, a poly-glycine linker or a gap was used to introduce multiple chains into AlphaFold (AF), a protein structure prediction program developed by Google DeepMind. AF recognized the linkers as unstructured regions and thus allowed to “dock” proteins and peptide chains to predict structures of the complex. Both methods have been benchmarked against protein-peptide and protein-protein complex datasets, achieving high accuracy in their predictions^6,7^. A surprising feature of the trained network is that no multiple sequence alignments (MSA) are needed for the peptide fragment when predicting protein-peptide complexes^6^. Consequently, a new AF neural network is now available, trained on oligomeric data for complex prediction. While its performance already surpasses state-of-the-art docking tools^7^, the weights of AlphaFold-multimer have not been extensively fine-tuned compared with AF and further improvements are expected.

Peptides mediate protein-protein interactions and often behave as a missing structural element that complements the structure of the receptor protein. Indeed, extended testing on AF for protein-peptide systems shows the ability to predict this type of interactions^6^. Peptides can be classified as interacting through short amino acid motifs (e.g. short turns), through secondary structure elements (e.g. hairpins and helices) or through discontinuous patches^8^. We chose to work on systems for which the interaction takes place through secondary structure elements as we believe this is the most complementary to the original AF training. Central to the success of AF, is its ability to assess the quality of the predictions through the predicted local-distance difference test (pLDDT) score. This per residue measure identifies regions of high and low accuracy in the predicted structure. However, when applied to complexes, there is no clear relationship between the pLDDT score and the binding affinity. Thus, peptides with very different binding affinity can be predicted with equally high pLDDT values.

The successes in predicting peptide-protein systems beg the question of whether AF can be used for peptide-protein screening of peptide libraries in a similar way that docking is used to filter small molecules in virtual screening. To be truly useful, we need some measure to rank order the quality of different predictions. We have previously used competitive binding simulations to predict relative binding affinities between peptides binding a certain protein receptor^9–11^. Though results have been able to distinguish strong, medium and weak binders, they can only be performed on a handful of systems due to computationally demanding simulations. In this work we expand the idea of competitive binding to AF predictions, in which two or more peptides are competing for the same binding site. Our results show that AF can indeed provide a way to screen thousands of sequences with relative ease (minutes per prediction) and accuracy. We show evidence of the robustness of this competitive binding methodology for several protein-peptide systems with experimentally characterized binding affinities. The method works best when the peptides bind along well-defined secondary structure elements and when the peptide sequences are not too similar to each other. A key requirement is that both peptide-protein complexes are well predicted in the binding site individually. We expect future improvements in AF trained complex neural network to provide higher ranking accuracy.

## Results

### The selected systems expand a wide range of binding affinities and binding motifs

We selected peptides that bind three different receptors, for which we had experimental binding affinities. First, the extraterminal domain (ET) of bromo and extraterminal domain proteins (BET) is an example of binding plasticity. Experimental structures of six different peptides binding ET reveal four different binding modes and a wide range of binding affinities, measured as K_d_, expanding the nano to micromolar range (see Fig. 1a and Table S1). The second and third receptors are MDM2 and MDMX, two proteins that bind an epitope of p53 as well as a series of possible inhibitors (see Fig.1a and Table S2). In this case, all peptides bind as a helix, with a wide range of binding affinities (reported as IC_50_s, see Table S2). The added benefit of this system is the difference of binding affinity of each peptide to MDM2 or MDMX, which we use as an example of competitive binding for the receptors.

**Figure 1.**
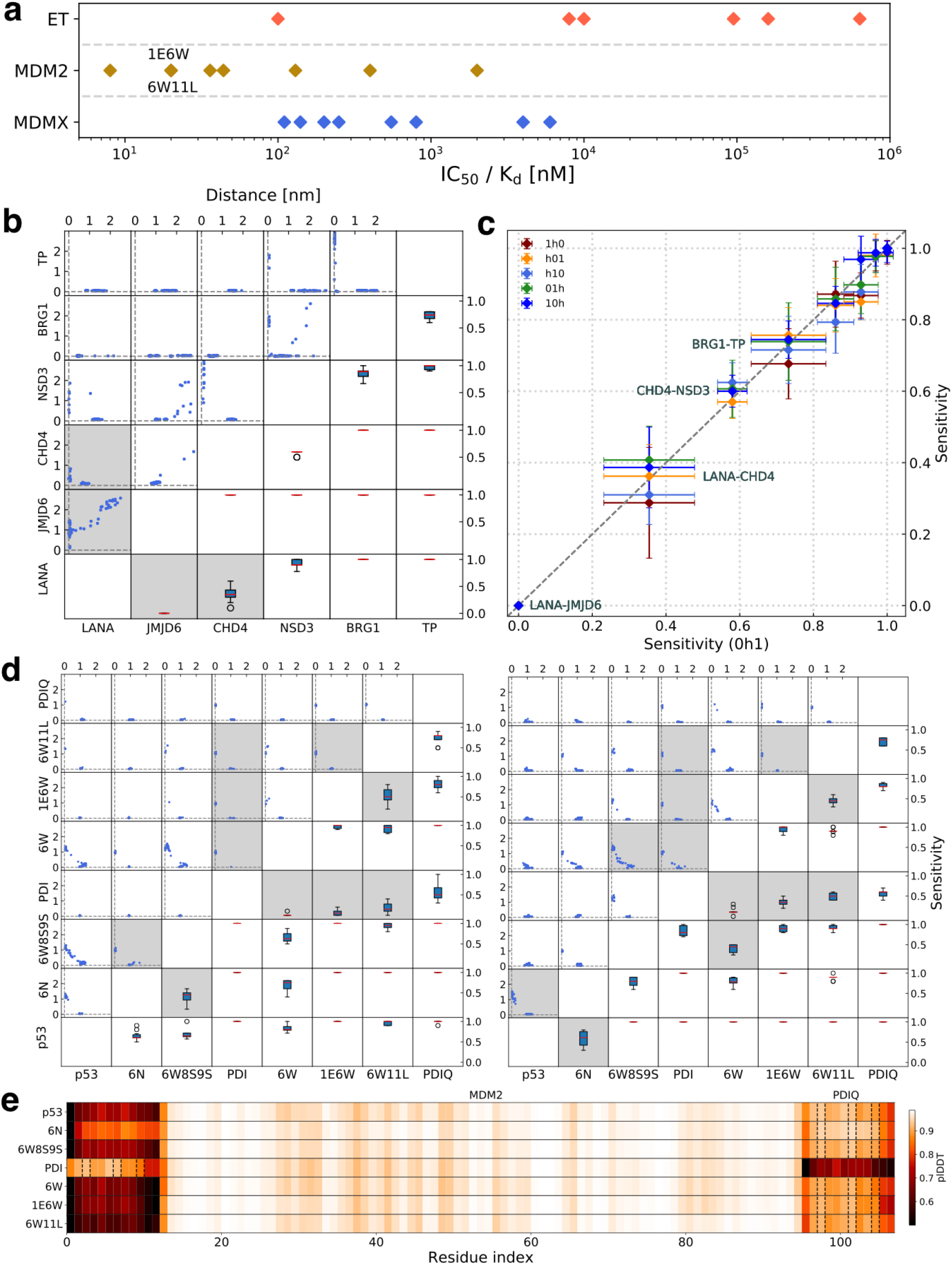
AF is capable of identifying high affinity peptide binders. **(a.)** The experimental K_d_ and IC_50_ values for ET and MDM2/MDMX-peptide systems, respectively. **(b.)** Prediction results for the ET domain with all pairs of peptides. Upper left grids (*g*_*i j*_, *i* + *j*≤ 4) show the distance of selected residues at the binding interface between binder at *x*_*i*_ versus *y* _*j*_ against their individual native structure from 100 predictions for each pair. Lower right grids (*g*_*i j*_, *i* + *j* ≥ 6) show the sensitivity value for each set of pairwise competitive binding test. Peptides are ordered by increasing binding affinity from left to right. Average sensitivity values lower than 0.5 are indicated by a gray block, indicating an AF competitive binding failure. **(c.)** Sensitivity comparison for different sequence inputs on ET-binders systems. For example, 0h1 stands for the input as peptide_0_:ET:peptide_1_. **(d.)** Prediction results for MDM2 (left) and MDMX (right) with selected pairs of peptides (increasing binding affinity from left to right). **(e.)** pLDDT values of the top rank model for PDIQ agasints all otehr peptides (binding MDM2). PDIQ was found at the binding pocket in all cases, except for PDI. The three hydrophobic anchoring residues are indicated by dotted lines.

Both ET and the MDM2/MDMX systems are important cancer targets, involving the folding upon binding of a peptide to the receptor protein. Competitive binding simulations are only meaningful if AF is able to predict accurate structures for each individual peptide/receptor pair. We first make predictions in which MSA is introduced for the protein receptor and no template or MSA is provided for the peptide. We find that AF was able to predict correct binding poses for all the MDM2/MDMX binding peptides. For the ET receptor, it failed to reproduce the experimental binding pose for one of the peptides (JMJD6)^11^ that binds as a helix. A more stringent test comes from providing no template or MSA for both the protein and peptide, which likely relies on AF having learned some biophysical principles during the training that allows it to identify good conformations^12^. While this approach is successful for predicting ET-peptide complexes, it fails to reproduce the binding pocket of MDM2/MDMX resulting in inaccurate complex predictions. (see Fig. S1)

### Competitive binding systematically predicts the three strongest binders to the ET domain

Peptides that interact with ET have an alternating hydrophobic/charged pattern of residues that allow a zipper like interaction with opposite charged residues in the receptor and a hydrophobic pocket. The remaining peptide sequence has low similarity between the different peptides (see Table S1) and dictates the secondary structure preferences upon binding (hairpin, single strand or helical). We performed AF competitive binding predictions for all possible pairings of the six known peptide binders, assessing the strongest binders by counting the number of times each peptide was found in the active site (see Figs. 1b, 2a and methods). For the three strongest binders (K_d_ of 0.1, 8 and 10 *µ*M), the results systematically show preference for the strongest binder (close to 100% of trials in the active site). However, along the weak binders the LANA peptide is predicted to be of similar binding affinity to CHD4 (K_d_ of 635 *µ*M and 95 *µ*M respectively) and a stronger binder than JMJD6 (K_d_=160 *µ*M). The case of JMJD6 was expected to be an outlier, as the bound state is not correctly predicted by AF. For the weak binding CHD4 and LANA, we find they both bind as a single strand, with few residues involved in direct interactions with the protein. The presence of alternating positively charged and hydrophobic residues leads to a registry shift strand pairing with the protein receptor in some predictions^11^. It is unclear how comparisons of peptides of different length will affect the rankings – we see here a case with CHD4 in which the longer LANA is preferred and the case of BRG1 of similar length to CHD4 which is correctly predicted as a stronger binder than both. As a sanity check, we show that the results are not dependent on the order in which the receptor and two peptides are introduced in AF (e.g. receptor-peptide_0_-peptide_1_ or peptide_1_-receptor-peptide_0_). (see Fig. 1c).

**Figure 2.**
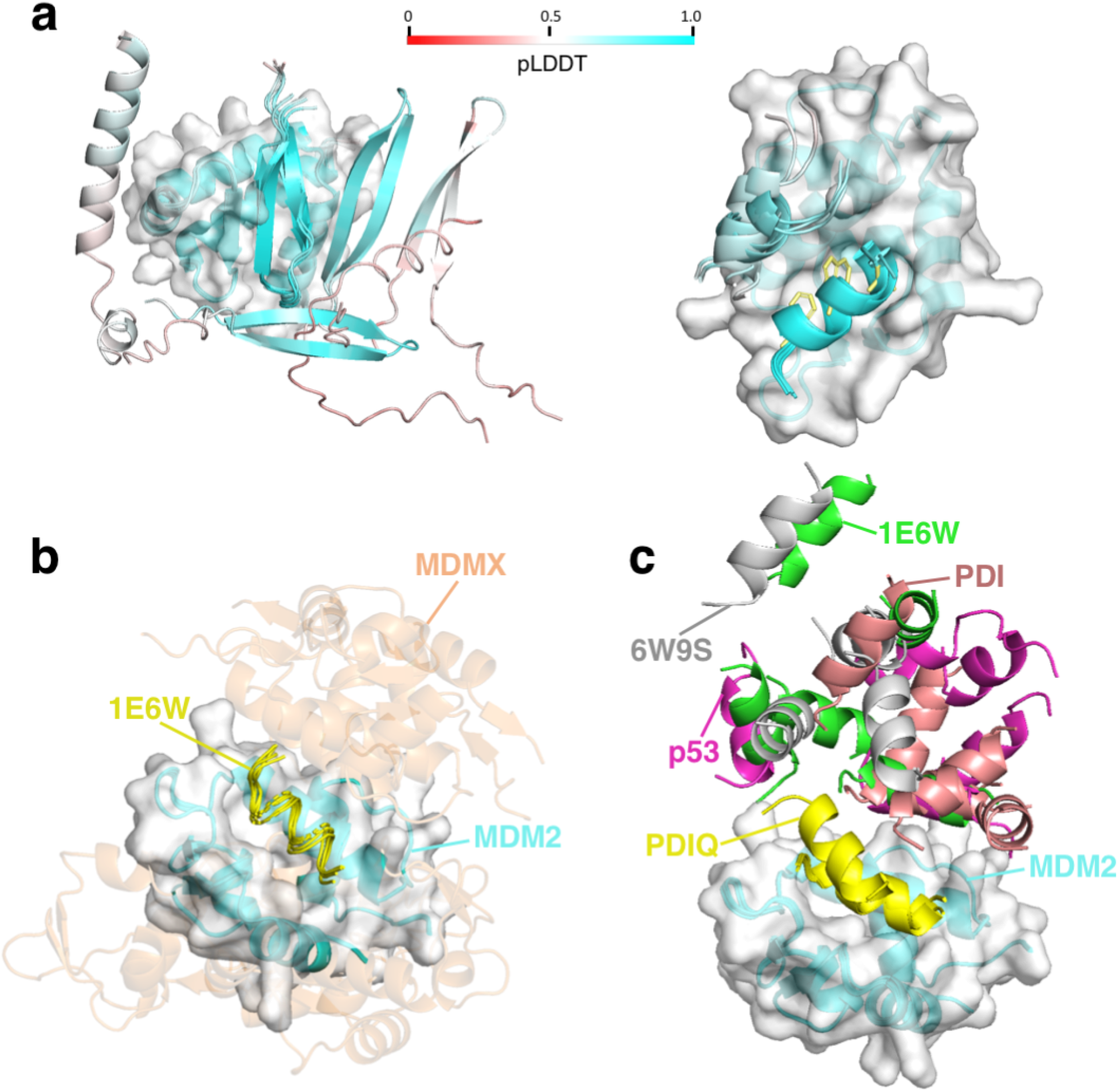
Prediction examples for screening high affinity binders via various AF based strategies. All receptor structures are aligned for visualization. **(a.)** Five prediction models from binary competitive binding for the ET domain (left) showing bound TP and unbound NSD3. For MDM2 (right) PDIQ is bound (highlighting three hydrophobic anchoring residues in yellow) and p53 is unbound. The color code for the peptides follows the pLDDT score, as shown in the colorbar. **(b.)** MDM2 (cyan) and MDMX (orange) compete for binding the peptide 1E6W (yellow). **(c.)** Five models from a multi-competitive binding prediction for MDM2 with 5 peptides (p53, 6W9S, PDI, 1E6W, PDIQ) consistently identify PDIQ as the strongest binder in the set.

### Low sequence variability in peptides binding MDM2/MDMX creates a greater ranking challenge

Binding to MDM2 and MDMX is characterized by three hydrophobic residues (Phenylalanine (F), Tryptophan (W), Leucine (L) at positions 3, 7 and 10, respectively) anchoring inside a hydrophobic pocket in MDM2 or MDMX (see Fig. 2a). As all peptides in the series are 12 residues long and contain these anchoring residues (see Table S2), we were uncertain of AF’s ability to capture binding differences. For example, AF predictions of three inactive peptides predict their binding to both MDM2 and MDMX. Furthermore, the sequence differences between some pairs of peptides is 2-3 residues, close to the limit of AF’s ability to differentiate as it was not parameterized for single point mutations^13^. Surprisingly, we find very good agreement between AF’s predictions and experimental ranking according to binding affinities (see Fig. 1d and S2). For MDM2, the PDIQ peptide is predicted to be the strongest binder for all pairings except one – the pairing with PDI gives a close to equal binding affinity to both, despite PDI being a weaker binder (8 nM versus 44 nM, see Table S2). Experimentally, all peptides are weaker binders to MDMX than MDM2, which reflect the challenges in developing dual inhibitors effective against MDMX. AF is not able to identify PMI (IC50=40nM) as the strongest binder in this set. Instead, PDIQ (the second-strongest binder for MDMX, IC50=110 nM) is selected as the strongest binder. Looking at the sequence similarity, the pMI peptide introduces a larger series of mutations than all other peptides in the series, which likely accounts for the incorrect ranking of this peptide in both MDM2 and MDMX. As with MDM2, the PDI peptide (IC_50_ = 550 nM) confounds AF, predicting it to be a binder of similar strength to PDIQ. We do not expect AF will be perfect in all cases, and these two peptides are good examples. However, the overall success in ranking the 11 peptides gives confidence in its value towards filtering down large peptide libraries.

As an additional sanity check, we competed each of the 11 peptides against itself to identify possible biases in the method. This competes peptides of equal binding affinity and sequence, and hence it is expected to be a toss coin (20 trials with 5 predictions in each). This is represented by a greater uncertainty in the results (see Fig. S3), where the average in most cases is close to 0.5. This is expected due to the small sampling size. Indeed, when we aggregate the results of the 11 trials (1100 predictions) the average is close to 0.5, with much smaller uncertainty (see last column in Fig. S3). Previous work using AF for peptide binding established that pLDDT score can be used to identify motifs in peptide-protein binding^6^. For competitive binding, we observe that the peptide at the binding interface generally shows higher pLDDT scores than the unbound one – even when single peptide AF predictions give high pLDDT scores for both (see Fig. 1e and S4).

### AF predicts peptides bind more tightly to MDM2 than MDMX

AF predicts accurate structures for individual peptide-MDM2 and peptide-MDMX simulations. Experimentally, the peptides bind with higher binding affinity to the MDM2 receptor. We introduced a new type of competitive binding in which one peptide competes for binding either receptor (MDM2 or MDMX). Interestingly, when providing MSAs for the receptors, AF invariably makes protein-protein complexes that disrupts the competitive binding assay. Such issues likely arise due to the co-evolution signals from MSAs of both receptors. Thus, we used the template feature of AlphaFold-multimer models for MDM2 and MDMX. No MSA is provided for either the peptide or the receptors (see Fig. 2b).

We found the peptide preferring the MDM2 pocket for 10 of the 11 peptides (see Fig. S5). The exception is pMI, which was already misbehaving in competitive peptide predictions and is again problematic for the receptor selection test. The binding affinity of pMI for the two receptors is similar (IC_50_ of 20 and 40 nM for MDM2 and MDMX respectively), with pMI being the strongest binder in the peptide series for MDMX while a top four for MDM2. As stated earlier, the greatest sequence variability in the series of peptides likely cause the insensitivity when competing with others. Despite this success, we noted some overlap between the MDM2 and MDMX structures, likely due to insufficient training of AlphaFold-multimer in the absence of MSAs as shown in Fig. 2b. We expect future iterations of AF to address such issues.

### Competitive binding is not successful for the binding of coils to MHC

Spurred by our current successes, we assessed the performance of the methodology on the Major Histocompatibility Complex Class II (MHC) as an important target for vaccine development. A series of 8 peptide sequences have been previously assessed with docking methodologies and their experimental binding affinities reported^14^. Unlike the previous examples, peptides bind MHC as an extended coil that interacts through a few amino acids. AF predicts binding at the correct site, but predicts higher helicity content than in the experimental structure (see Fig. S6a). In contrast with previous systems, AF has very low confidence on the peptide structures, as indicated by very low pLDDT scores (see Fig. S6b). Not surprisingly, competitive binding modeling fail in this scenario (see Fig. S6c).

### Pushing beyond pairwise binder selection and current limitations

Finally, we pushed the method to its limit for computational efficiency. Could we compete more than 2 peptides? We find that as more and more peptides were added, AF has a tendency to create protein assemblies from the different peptide fragments to mimic protein structures (see Fig. S7c). We observed assemblies for peptides that have a tendency to bind as strands (e.g. binding the ET domain). While for those that bind as helices (MDM2/MDMX) we identified the strongest binder in the active site (see Fig. 2c). Given that each prediction run takes a few minutes on an A100 GPU, we see no real advantage in competing more than 2 peptides.

## Discussion

AF has already surpassed the ability of other docking programs in predicting peptide-protein complexes, according to independent studies. While there are still limitations, AF gives the opportunity of screening large libraries of peptide-protein complexes. Peptide epitopes are typically mediating protein-protein interactions, and many such epitopes can be identified from bioinformatics or experimental pull-down experiments. However, experimental validation is typically slower, as processing the hundreds of possible hits will require extensive resources. AF screening can be a useful tool to discern those sequences that are likely to bind. And using AF in competitive binding mode allows identifying those peptides that might have the highest binding affinity. This application works best when we have a few peptides of known binding affinity. This can be instrumental for structure determination. For example, NMR determination of protein-peptide complexes using NOESY will suffer from broad peaks with weak binders, resulting in a costly and lengthy structure determination process. On the other hand, strong binders (at the same concentration) will result in sharper NOESY peaks that lead to fast and accurate structure determination of the structures.

There are some expected limitations of the methodology. We first expect the methodology to work better for peptides that adopt well-defined secondary structures upon binding. This idea equates to thinking about the peptide binder as a missing part of the protein. For competitive binding simulations to work, AF must first be able to predict the correct bound structures for each peptide independently with high confidence (e.g. high pLDDT score). We find this is not the case for the JMJD6-ET interaction, and indeed the correct ranking between the LANA and JMJD6 peptide is incorrectly determined. Using the protocols described here, users should first check the single peptide-protein prediction of known binders for their system of interest, before attempting competitive binding runs. Finally, just as AF is not sensitive to single point mutations, competitive binding for peptides with very similar sequences will not be efficient for current models.

A recent work^12^ suggests that AF’s MSA allow focusing conformational search near the global energy minima, and the trained AF model is able to refine the structures based on a learned potential energy. We find that indeed, providing no MSA for the protein receptor might work in some cases (see Fig. S7a), but not always (see Fig. S7b). On the other hand, providing either a template or a MSA for the receptor will allow AF to identify the correct structure of the receptor and use its learned potential to identify the peptide’s binding site and conformation. In the cases where a peptide competes for binding two or more receptors, using MSA will lead to AF docking the two receptors – thus, in these cases templates are recommended if available. Using AF to rank order peptides according to binding affinities seems to work specially well when competing strong binders against weak binders. We do not expect this method to work in all cases, and indeed we find a few exceptions. However, it’s robustness and current success make it a valuable tool for sorting through libraries of possible binders. It can be specially useful when a few peptides already have experimentally determined binding affinities. As expected, comparing peptides of similar binding affinity showed greater variability and increased uncertainty. Competing weak binders can sometimes result in predictions where neither peptide is observed in the binding site. We expect that as the training of AF for complexes improves, and with further improvements in machine learning models, the approach we have outlined here will result in higher accuracy predictions and ranking of peptide sequences. Given that some protein-protein interfaces are mediated by peptide epitopes in coil conformation, future advances that accurately predict such conformations will be invaluable towards screening of a larger fraction of systems (e.g. the MHC).

## Methods

### AlphaFold competitive peptide binding predictions

Modeling was performed on a local install of the publicly available ColabFold (https://github.com/sokrypton/ColabFold)^15^. MSA search was done by MM-seqs2^16^ on UniRef100 database^17^. For the input we use gaps to separate different chains (e.g. for the competitive binding of p53 and PDIQ to MDM2: ETFEHWWSQLLS:…:ETFSDLWKLLPE, where the sequence of MDM2 was omitted) and allow MSA on the protein receptor, but only query the sequence itself for both peptide sequences. AF monomer models and no template were used for prediction unless otherwise noted. For each binary competitive modeling, we ran AF predictions 20 times (5 predictions each) to accumulate 100 predictions. Each run takes ∼4 minutes on an A100 GPU. For each prediction, we determined the receptor-peptide center **c**_**1**_ (for peptide 1) and **c**_**2**_ (for peptide 2) by averaging the coordinates of selected residues from receptor and peptide and compare with the center in the native structures for each peptide 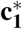 and 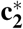 by 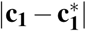 versus 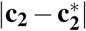. Table S3 summarizes the residue selections for both peptide binders and receptors for the distance calculations. Typically, one of the peptides will match the experimental structure and the other be far away from the binding site. We randomly split all 20 prediction runs into 10 sets to calculate the sensitivity for each set as 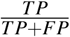, where *TP* and *FP* stands for true positive and false positive, respectively. Uncertainty was estimated from the sensitivity values of all 10 sets.

## Acknowledgements

We are thankful for the use of Hipergator computational resources at the University of Florida.

## Author contributions statement

L.C and A.P conceived and planned the research, L.C. conducted experiments and analyzed the results, L.C. and A.P. discussed the results, drafted and reviewed the manuscript.

