## Supplemental tables and figures for "AlphaFold encodes the principles to identify high affinity peptide binders"

**Table 1.** $K_d$ values for peptides bound to ET domain sorted from lowest to highest affinity.

| Name | Sequence | $K_d$ (μM) |
|---|---|---|
| LANA | NLQSSIVKFKKPLPLTQPG | 635[1] |
| JMJD6 | KWTLERLKRKYRN | 160[2] |
| CHD4 | KVAPLKIKLGGF | 95[3] |
| NSD3 | PEIKLKITKTIQNGRELFESSLCGDLLNEVQASE | 10[4] |
| BRG1 | RSVKVKIKLGRK | 8[3] |
| TP | SRLTWRVQRSQNPLKIRLTREAP | 0.1[4] |

**Table 2.** $IC_{50}$ values for peptides bound to MDM2/X sorted from lowest to highest affinity for MDM2.[5]

| Name | Sequence | MDM2 (nM) | MDMX (nM) |
|---|---|---|---|
| p53 | ETFSDLWKLLPE | 2000 | 6000 |
| 6N | LTFEHNWAQLTS | 400 | 4000 |
| 6W8S9S | LTFEHWWSSLTS | 130 | 800 |
| 6W9S | LTFEHWWASLTS | 125 | 500 |
| PDI | LTFEHYWAQLTS | 44 | 550 |
| 6W | LTFEHWWAQLTS | 36 | 250 |
| 6W8S | LTFEHWWSQLTS | 24 | 180 |
| 1E6W | ETFEHWWAQLTS | 20 | 200 |
| 6W11L | LTFEHWWAQLLS | 20 | 140 |
| pMI | TSFAEYWNLLSP | 20 | 40 |
| PDIQ | ETFEHWWSQLLS | 8 | 110 |
| 6S | LTFEHSWAQLTS | Inactive | Inactive |
| 1E6N | ETFEHNWAQLTS | Inactive | Inactive |
| 4T6W | LTFTHWWAQLTS | Inactive | Inactive |

**Table 3.** Residue selection for binding interface distance calculation between predicted and native structure, all residues were selected for molecules not shown.

| Name | Selected residues |
|---|---|
| ET | 39 - 43 |
| LANA | 5 - 9 |
| CHD4 | 6 - 7 |
| NSD3 | 2 - 7 |
| BRG1 | 3 - 8 |
| TP | 13 - 19 |
| MDM2 | 30-37, 46-47, 67-69 |
| MDMX | 30-40, 49-50, 70-73 |

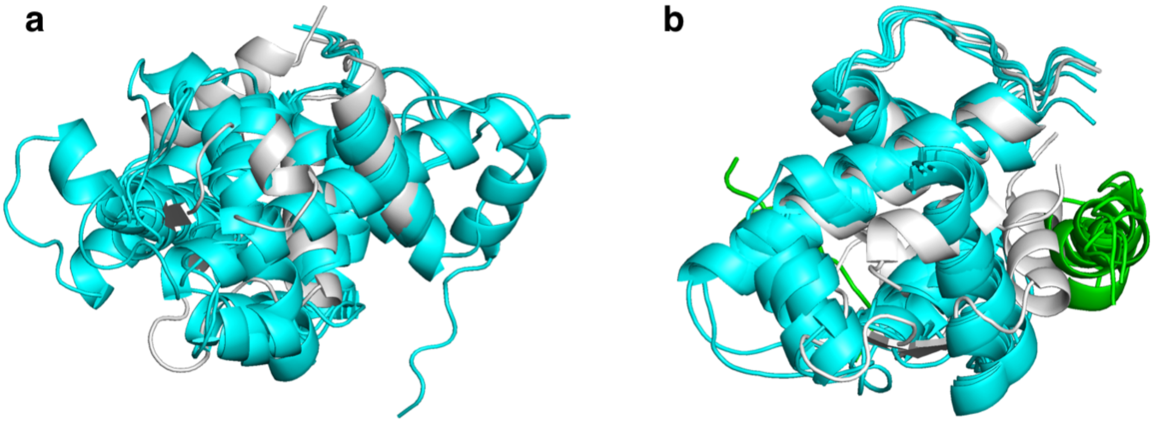


**Supplementary Figure 1. AF predictions in the absence of templates or MSA for MDM2 fail to identify the native structure. (a.)** AF predictions of MDM2 (cyan) aligned with the native structure (white). **(b.)** AF predictions of the MDM2-p53 complex (cyan for MDM2 and green for p53) aligned with the native structure (white).

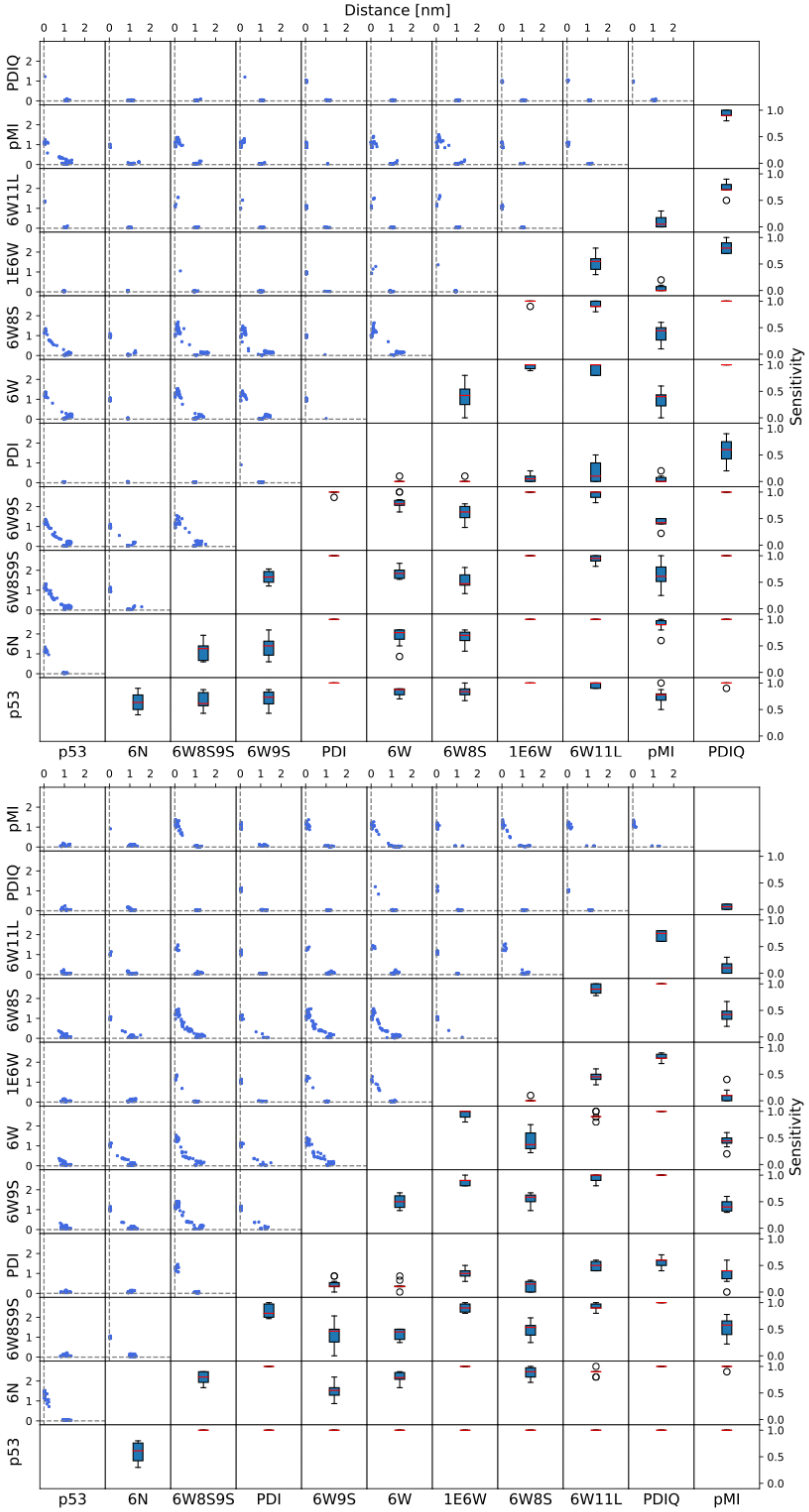


**Supplementary Figure 2. Prediction results for competitive binding of all peptide pairs with MDM2 (top) and MDMX (bottom).** Upper left grids ($g_{ij},\ i+j \leq 9$) show the distance of selected residues at the binding interface between binder at $x_i$ versus $y_j$ against their individual native structure from 100 predictions in total for each pair. Lower right grids ($g_{ij},\ i+j \geq 11$) show the sensitivity value for each set of pairwise competitive binding test.

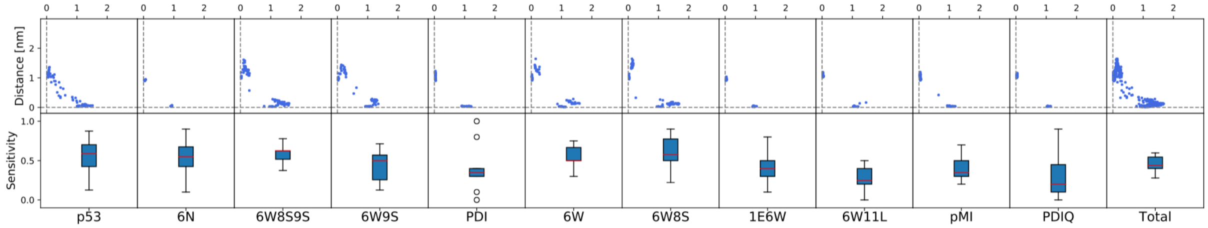


**Supplementary Figure 3. Baseline prediction results for competitive binding of peptides with MDM2.** Sequence input for baseline test is MDM2 with duplicate peptides (e.g.: p53:p53:MDM2). Row 1: distance between predicted bound structure with experimental native structure at the center of binding interface for each peptide; row 2: sensitivity of identifying receptor bound with either binder for each peptide. The last column shows an aggregate of all prediction results. The average and uncertainty were evaluated based on 11 peptides, with 100 predictions for each.

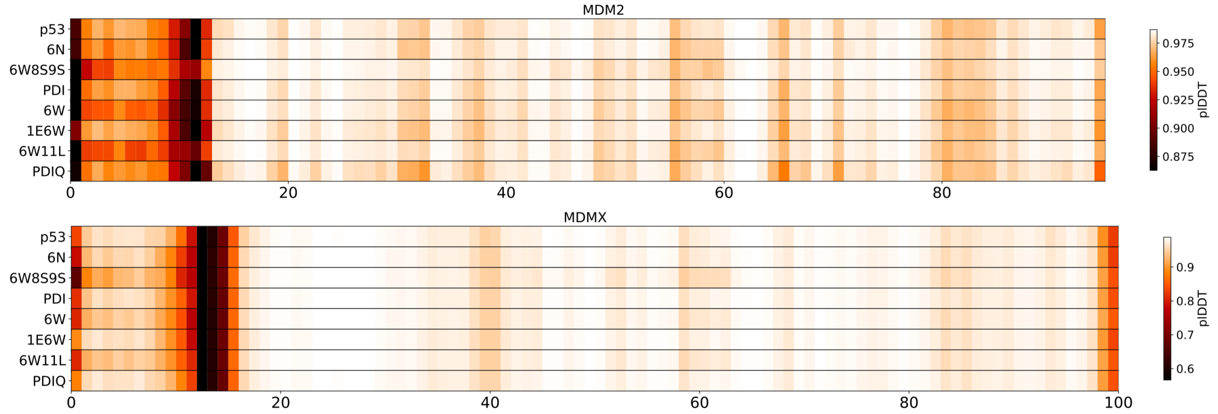


**Supplementary Figure 4. pLDDT values for the top ranked prediction for a single peptide bound to MDM2 (top) or MDMX (bottom).** Input of single peptide and receptor was separated by ":" gap with MSA for the receptor. Predictions were done with AF monomer models.

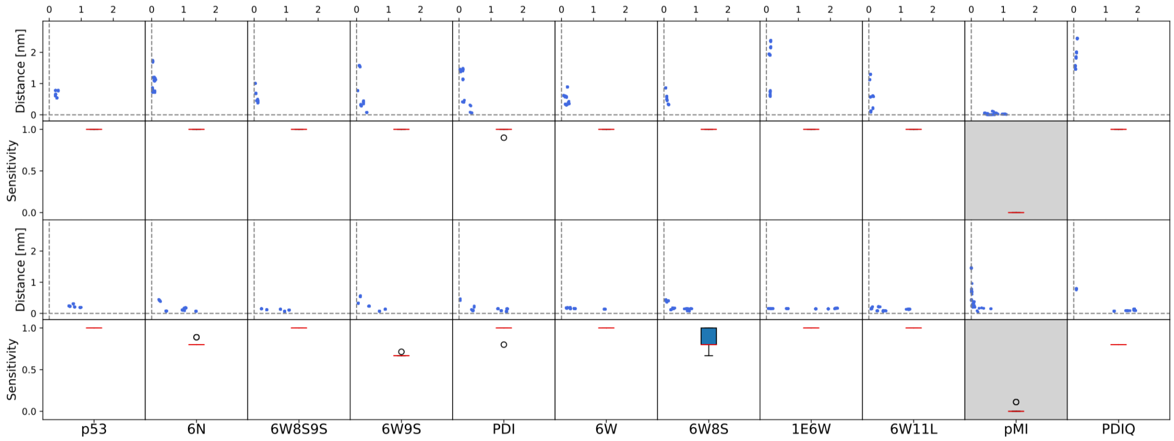


**Supplementary Figure 5. Prediction results for competitive binding of single peptides against MDM2 and MDMX.** For Rows 1 and 2 the AF input was: MDM2:peptide:MDMX; For Rows 3 and 4 the input was MDMX:peptide:MDM2. Rows 1 and 3 show the distance between the center of binding interface for each peptide and each of the receptors. Rows 2 and 4 report the sensitivity of identifying the peptide bound to MDM2 or MDMX. Both sets of predictions favor binding MDM2, except pMI which favors binding to MDMX.

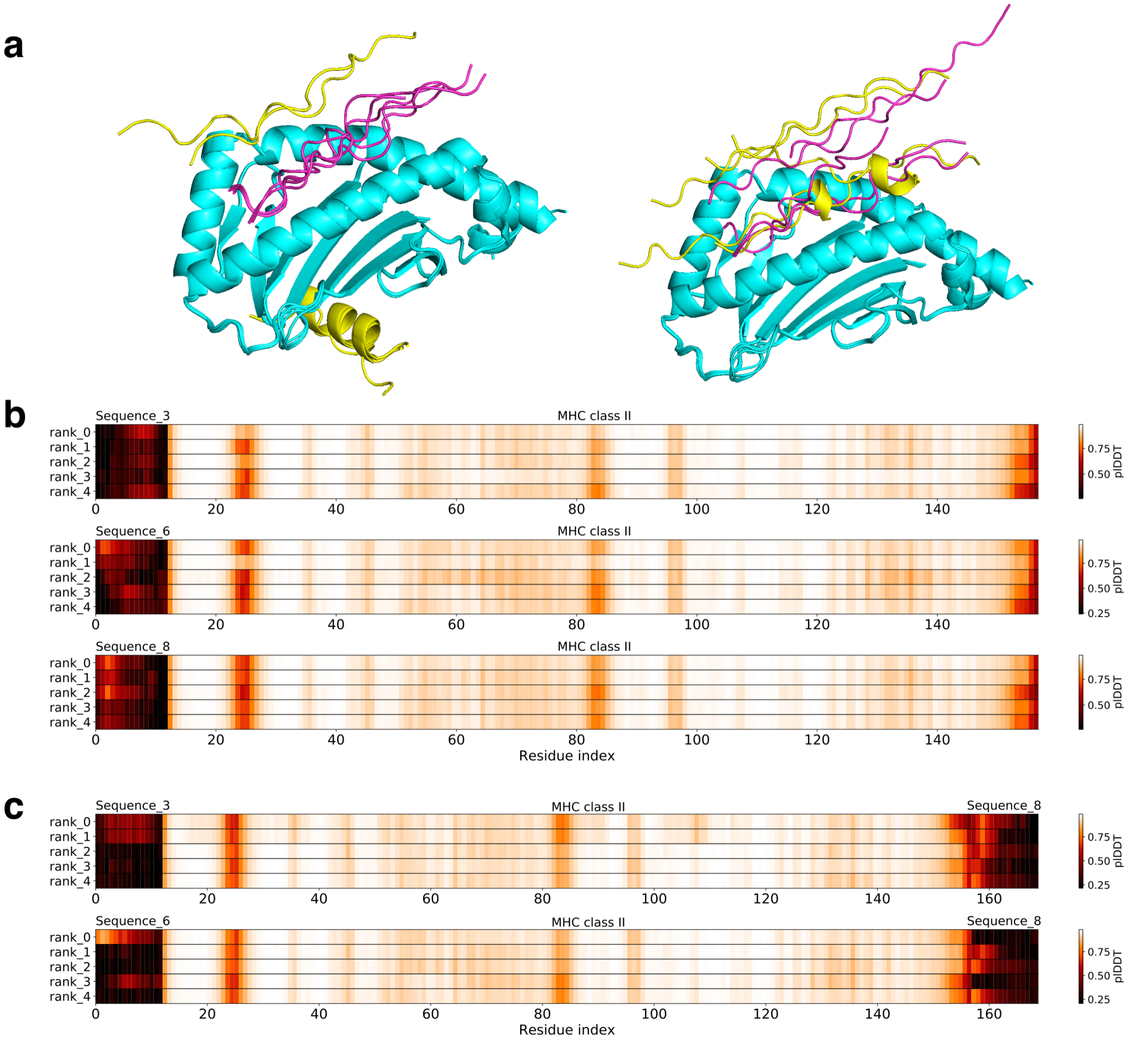


**Supplementary Figure 6. Prediction results for competitive binding of selected peptide pairs with the MHC class II receptor. (a.)** Five output models aligned on the receptor showing competitive binding of sequence 3 (yellow) and 8 (purple) on the (left) and sequence 6 (yellow)) vs 8 (purple) (right). **(b.)** pLDDT scores of single peptide bound to receptor for sequence 3, 6, 8. **(c.)** pLDDT scores of binary competitive binding for sequences 3 vs 8 (top) and sequences 6 vs 8 (bottom). The sequence index followed the affinity rank from Ochoa *et al.*[6].

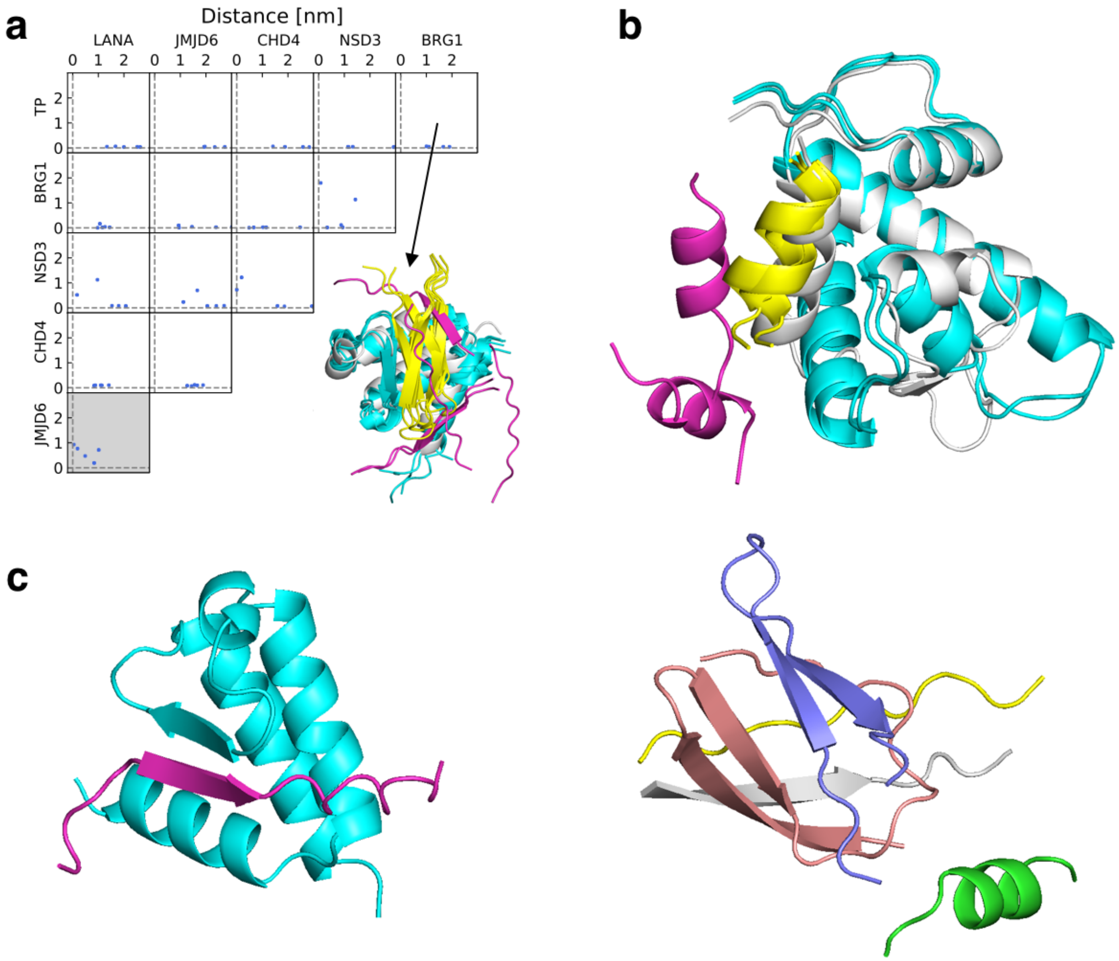


**Supplementary Figure 7. Single sequence prediction results for binary competitive binding and self assembly example from multi-competitive binding. (a.)** (top left) Each grid shows the distance of selected residues at the binding interface between binder at $x_i$ versus $y_j$ against their individual native structure from 5 predictions without MSA or template for each pair. (bottom right) Predicted structures for BRG1 (purple) and TP (yellow) competitive binding with ET (cyan) and the native structure of ET (white). **(b.)** Predicted structures without MSA or template for p53 (purple) and PDIQ (yellow) competitive binding against MDM2 (cyan) and the native structure of MDM2 (white). **(c.)** Selected output model for multi-peptide competitive binding against ET (cyan). One peptide is found bound (purple) where the other five peptides assemble far from the protein receptor.
